# Molecular modeling predicts novel antibody escape mutations in respiratory syncytial virus fusion glycoprotein

**DOI:** 10.1101/2022.02.25.482063

**Authors:** Sierra S. Beach, McKenna Hull, F. Marty Ytreberg, Jagdish Suresh Patel, Tanya A. Miura

## Abstract

Monoclonal antibodies are increasingly used for the prevention and/or treatment of viral infections. One caveat of their use is the ability of viruses to evolve resistance to antibody binding and neutralization. Computational strategies to predict which mutations will result in antibody resistance would be invaluable because current methods for identifying potential escape mutations are labor intensive and system-biased. Respiratory syncytial virus is an important pathogen for which monoclonal antibodies against the fusion (F) protein are used to prevent severe disease in high-risk infants. In this study, we used an approach that combines molecular dynamics simulations with FoldX to estimate changes in free energy in F protein folding and binding to the motavizumab antibody upon each possible amino acid change. We systematically selected 8 predicted escape mutations and tested them in an infectious clone. Consistent with our F protein stability predictions, replication-effective viruses were observed for each selected mutation. Six of the eight variants showed increased resistance to neutralization by motavizumab. Flow cytometry was used to validate the estimated (model-predicted) effects on antibody binding to F. Using surface plasmon resonance, we determined that changes in the on-rate of motavizumab binding were responsible for the reduced affinity for two novel escape mutations. Our study empirically validates the accuracy of our molecular modeling approach and emphasizes the role of biophysical protein modeling in predicting viral resistance to antibody-based therapeutics that can be used to monitor the emergence of resistant viruses and to design improved therapeutic antibodies.

**Importance:** Respiratory syncytial virus (RSV) causes severe disease in young infants, particularly those with heart or lung diseases or born prematurely. As no vaccine is currently available, monoclonal antibodies are used to prevent severe RSV disease in high-risk infants. While it is known that RSV evolves to avoid recognition by antibodies, screening tools that can predict which changes to the virus will lead to antibody resistance are greatly needed.

## Introduction

The use of monoclonal antibodies (mAb) for the treatment or prevention of viral infections has accelerated over the past few years. mAb are a powerful tool against viral infections because of the ability to target specific epitopes to neutralize viral pathogens. The first mAb approved by the FDA for the prophylaxis of viral infection was Synagis® (palivizumab), which is used to prevent respiratory syncytial virus (RSV) infection in high-risk infants (1–3). One of the most notable recent uses of mAb has been the emergency use authorization of three mAb treatments for severe acute respiratory syndrome coronavirus 2 (SARS-CoV-2) the causative agent for coronavirus disease 2019 (COVID-19) (4–7). The FDA has also approved the mAb cocktail, Inmazeb™, for the treatment of Ebola virus infection (8) and Trogarzo®, a mAb treatment for drug-resistant human immunodeficiency virus (HIV-1) infection (9). There are also mAb therapies in development for other viruses including influenza virus (10) and herpes simplex virus 1 (11, 12). The use of mAb for the prevention and/or treatment of viral infections has become an important strategy for the medical community.

One caveat of using mAb against viral infections is the increased selective pressure for antibody escape mutations to occur. The current methodology for monoclonal antibody-resistant mutants (MARMs) discovery involves sequencing samples taken from patients or serially passaging virus with the antibody of interest to select for evolved mutations. This presents challenges in that the escape mutation is already circulating in the population and passaging experiments can be time consuming and are biased in only detecting mutations that can arise over limited replication time in cell culture. Given that it is now common to monitor the genetic variation in a population, an aspirational goal in the fight against infectious disease is the ability to predict when viral evolution is about to outpace the effectiveness of protective mAb.

Paramount to this goal is the ability to predict amino acid changes that will significantly disrupt binding of antibodies to viral proteins; a collection of these predicted MARMs would constitute a watch list. An increase in the frequency of MARMs on the watch list in an infected population could signal reduced effectiveness of antibody-based therapies and the possible adaptation to these therapies.

Protein biophysical models can be used to predict protein stability and the disruption of interactions with other biomolecules due to mutations. In our previous studies, we used FoldX software combined with molecular dynamics (MD) simulations (MD+FoldX) to estimate folding and binding stabilities of Ebola virus (EBOV) envelope glycoprotein (GP) and mAbs KZ52, Antibody 100 (Ab100), Antibody 114 (Ab114) and 13F6-1-2. Using this approach, we generated a watch list comprised of 127 mutations that were predicted to disrupt binding between GP and four mAbs but would not disrupt the ability of the glycoprotein to fold and form a trimer (13–16). Three mutations from our watch list have already been seen in humans or are experimentally known to reduce efficacy of the antibody treatment (17, 18). While these previous studies showed the predictive capability of protein biophysical modeling to help guide empirical science in examining phenotypes in viral evolution, these results were not empirically validated to test the accuracy in predicting MARMs.

MARMs for palivizumab have been identified from cell culture, animal models, and clinical samples (19–27). RSV is an important pathogen for infants, elderly, and immune compromised that causes severe lower respiratory infections and is the second leading cause of infant death in the world (2, 3). The only targeted treatment for RSV is the prophylactic mAb palivizumab and additional mAbs are in development, including nirsevimab, which is currently in Phase 3 clinical trials (28–32). Palivizumab and its derivative, motavizumab, target the site II epitope on the fusion glycoprotein (F protein), the protein responsible for the fusion of viral and host cell membranes during viral entry (28, 33–38). F protein is a desirable target for anti-viral treatments given the function of the protein for viral entry, the response to the antigen by the host immune system, and the conservation of antigenic sites including site II among RSV strains (38–43). F protein is also advantageous to use to test molecular modeling given the multiple co-crystal structures with mAbs (44–47). For this study we used the co-crystal structure of motavizumab-F protein because there is no crystal structure with palivizumab. Motavizumab is also of interest as only one MARM, K272E, has ever been derived from clinical and empirical lab studies (21).

In this study we describe the application and empirical validation of our MD+FoldX approach to RSV F protein and motavizumab complex to predict MARMs by analyzing changes in relative binding affinities due to all possible amino acid changes in the F protein. We empirically tested the fitness, mAb neutralization, and binding characteristics of eight predicted MARMs using an infectious clone. Our modeling approach was able to predict the already known MARM, K272E, and five additional novel variants with decreased neutralization by motavizumab.

## Results

### Molecular modeling identified amino acid residues of interest in F monomer and motavizumab interaction

MD+FoldX approach was used to predict RSV F protein escape mutations that would allow the F protein to fold correctly but disrupt its recognition by motavizumab. We estimated the effect of all possible mutations of RSV F protein for all 448 amino acids and 19 possible substitutions at each site for both F protein monomer folding (ΔΔ*G*_Fold_) and its binding with motavizumab (ΔΔ*G*_Bind_) (Figure 1 and Appendix). Mutations with ΔΔ*G*_Fold_ value less than 2 kcal/mol are not predicted to affect the stability of F protein and therefore have the potential to arise under selective pressure (15). To identify MARMs and test the accuracy of the MD+FoldX approach, we selected eight mutations with ΔΔ*G*_Bind_ values ranging from 0.5 to 5.5 kcal/mol (low to high disruption of binding to motavizumab) and ΔΔ*G*_Fold_ values less than 2 kcal/mol (Figure 1, Table 1). K272E, which had a ΔΔ*G*_Bind_ value of approximately 2 kcal/mol and a ΔΔ*G*_Fold_ value less than 0 kcal/mol, was included as a positive control as the only known escape variant for motavizumab (21).

**Table 1.**
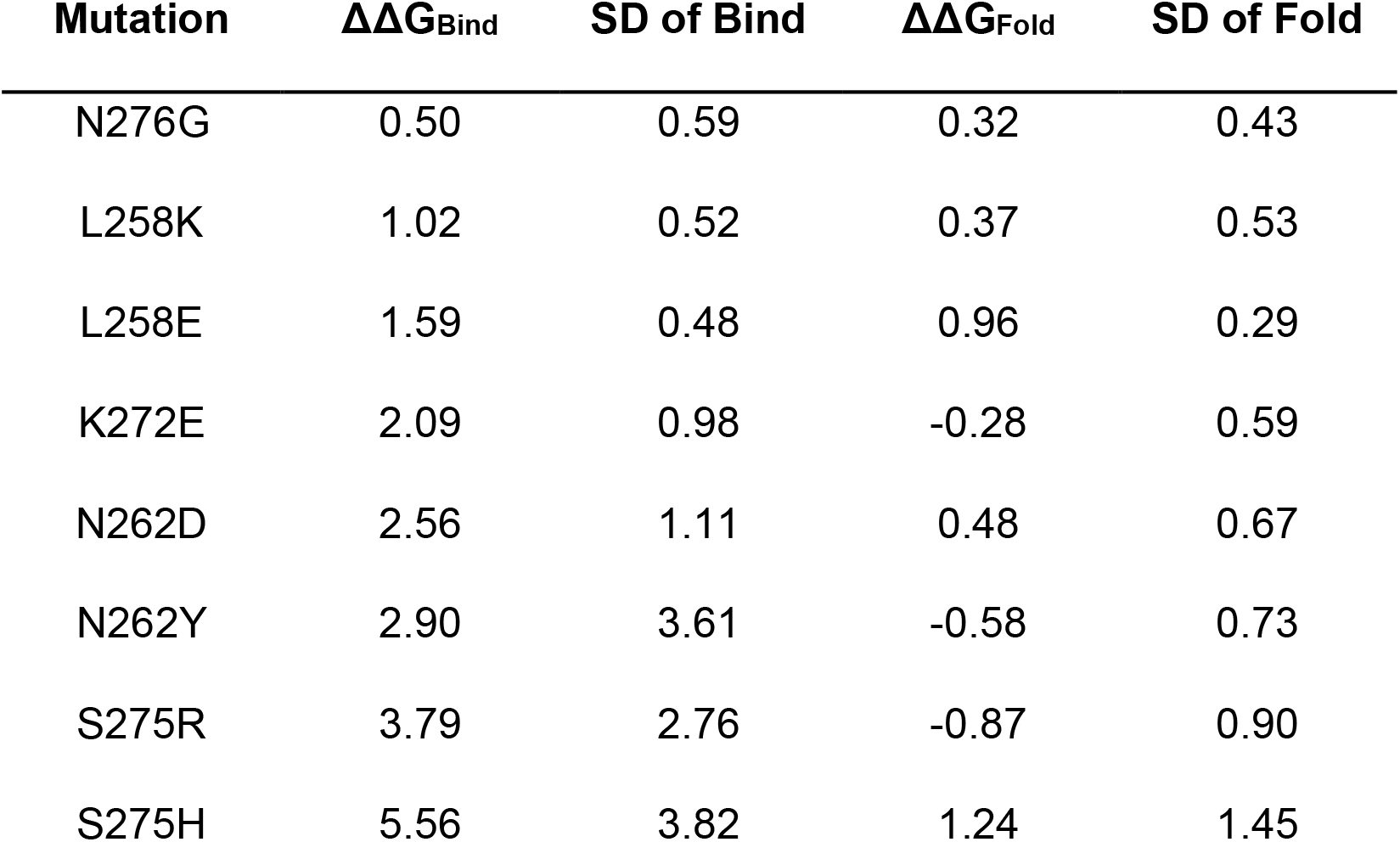
ΔΔ*G*_Fold_ and ΔΔ*G*_Bind_ values and standard deviations of eight selected variants.

| **Mutation** | **$\Delta\Delta G_{\text{Bind}}$** | **SD of Bind** | **$\Delta\Delta G_{\text{Fold}}$** | **SD of Fold** |
| --- | --- | --- | --- | --- |
| N276G | 0.50 | 0.59 | 0.32 | 0.43 |
| L258K | 1.02 | 0.52 | 0.37 | 0.53 |
| L258E | 1.59 | 0.48 | 0.96 | 0.29 |
| K272E | 2.09 | 0.98 | -0.28 | 0.59 |
| N262D | 2.56 | 1.11 | 0.48 | 0.67 |
| N262Y | 2.90 | 3.61 | -0.58 | 0.73 |
| S275R | 3.79 | 2.76 | -0.87 | 0.90 |
| S275H | 5.56 | 3.82 | 1.24 | 1.45 |

**Figure 1.**
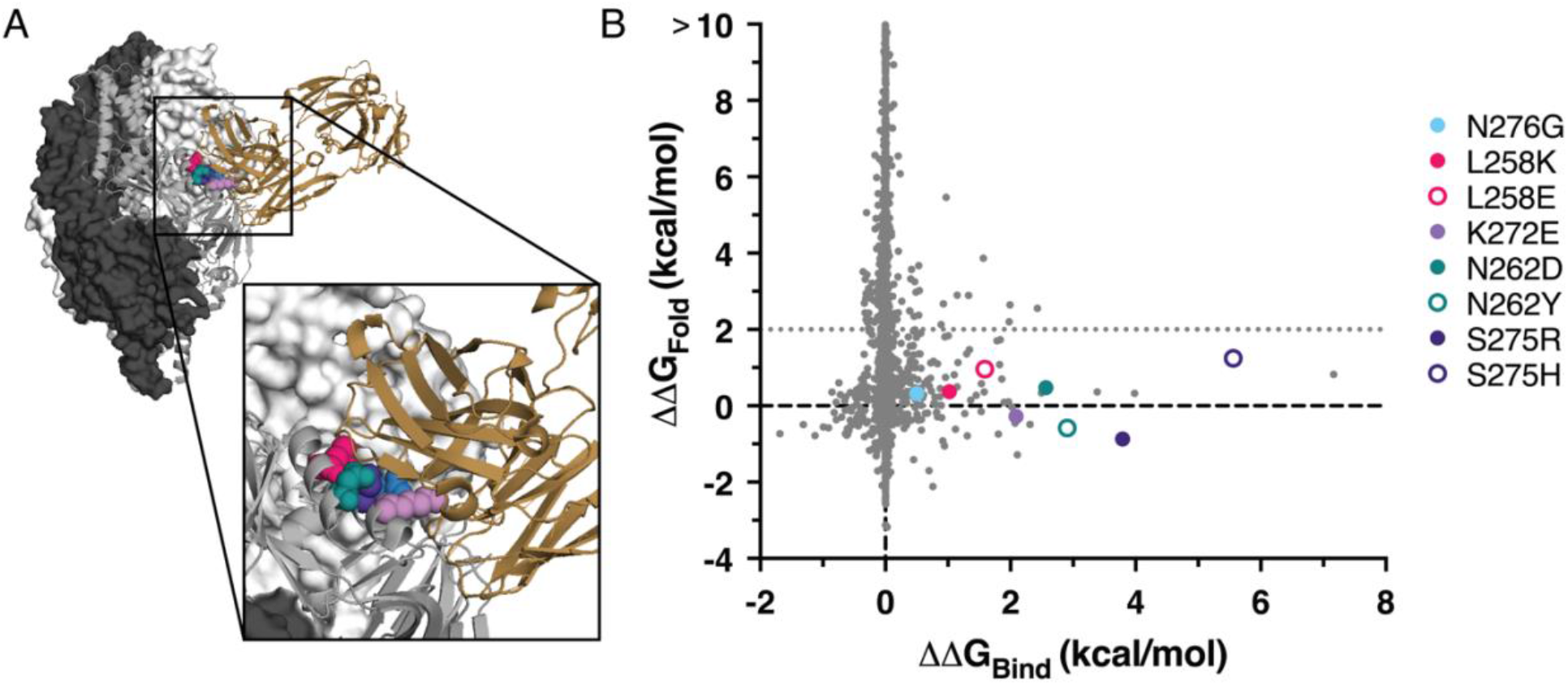
Molecular modeling of F protein and motavizumab interaction. (A) F protein trimer (dark gray) monomer (gray ribbon) with motavizumab (gold ribbon) structure with interacting residues of interest: L258 (pink), N262 (teal), K272 (lavender), S275 (purple), and N276 (blue). Based on PDB ID:4ZYP. (B) Plot of MD+FoldX predictions of ΔΔ*G*_Fold_ and ΔΔ*G*_Bind_ of all possible mutations in F with eight selected variants.

### The eight selected variants were viable and demonstrated different growth kinetics

To confirm that ΔΔ*G*_Fold_ values less than 2 kcal/mol did not disrupt F protein function, we first assessed the ability of the eight variants to replicate. To test the growth kinetics of the variants, site directed mutagenesis was used to create the variants, which were subcloned into a recombinant mKate2-expressing RSV infectious clone (48). HEp-2 cells were infected at a multiplicity of infection (MOI) of 0.1 and supernatant media samples were collected over 48 hours then titrated by TCID_50_ assay. We were able to generate replicating virus from each mutated clone (Figure 2). L258K and L258E both demonstrated delayed growth with extracellular virus remaining undetectable until 15 and 18 hours, respectively. These variants also exhibited significantly reduced overall viral titers when compared to wild-type (WT) over the 48-hour period. N262D and N262Y also displayed significantly reduced viral titers when compared to WT. S275R and S275H were similar to WT virus in growth with small differences in titers at a few of the time points. K272E had reduced viral titers at each time point and N276G had few differences from WT. All eight mutations were able to replicate but displayed at least some reduced propagation in viral titer when compared to the WT virus.

**Figure 2.**
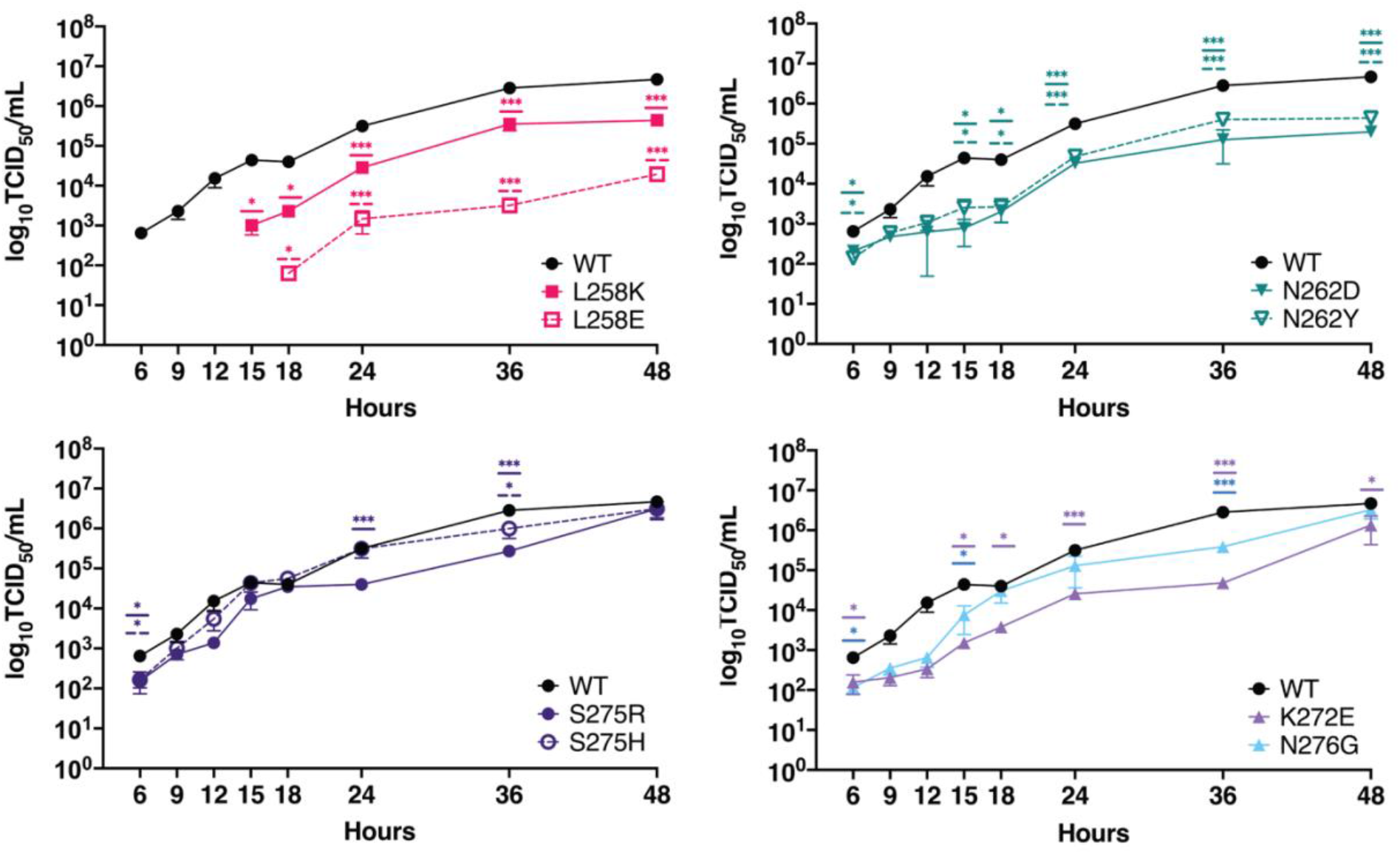
All eight variants demonstrated ability to replicate and propagate new virus. HEp-2 cells were infected with 0.1 MOI of variant or WT virus and supernatant media were collected over time. Virus was titrated by TCID_50_ assay on HEp-2 cells and is shown as the average and SEM of three independent experiments. \**p* <0.05; \*\**p*<0.005; \*\*\**p*<0.001.

### Six variants demonstrated decreased neutralization by motavizumab compared to WT

After establishing that all eight variants were able to replicate the next step was to quantify neutralization of the variants by motavizumab. Virus variants were incubated with a two-fold serial dilution of motavizumab for 1 hour prior to infection of HEp-2 cells and mKate2-expressing cells were quantified by flow cytometry after 18 hours. Six of the eight mutations demonstrated reduced neutralization by motavizumab compared to WT (Figure 3). The only known MARM, K272E, was not neutralized at the highest dose of antibody used (10 μg/mL). Previously published data determined that the IC_50_ for K272E was 30.04 ± 611.35 μg/mL (21). S275H, L258K, and S275R demonstrated a greater than 5-fold increase of IC_50_ when compared to WT and L258E and N262D had a greater than 3-fold increase in IC_50_ (Figure 3, Table 2). Variants N262Y and N276G had similar neutralization kinetics to WT (Figure 3, Table 2). In addition to confirming K272E as a motavizumab-resistant mutation, we identified 5 novel MARMs that have not been previously identified by traditional approaches.

**Table 2.**
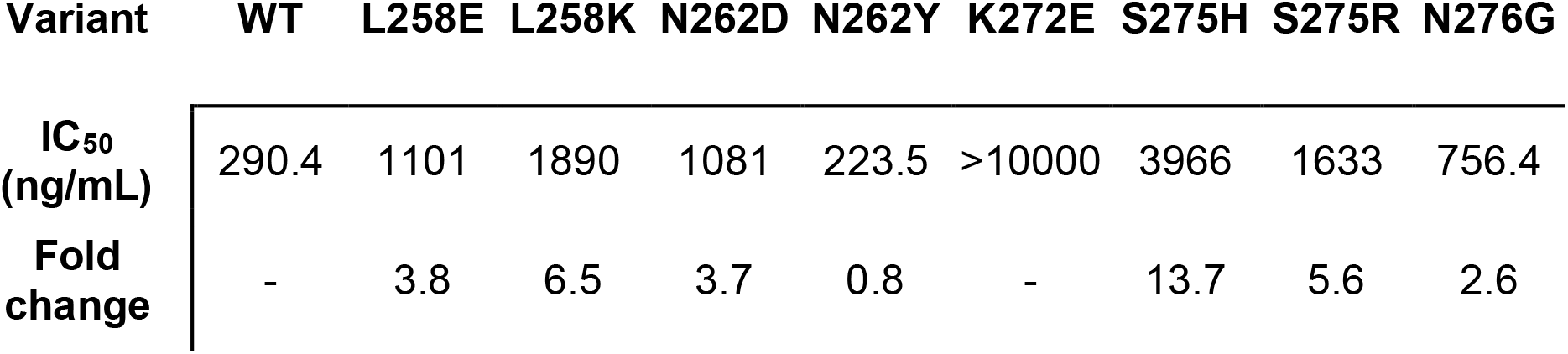
IC_50_ values and fold change compared to WT of selected variants.

| Variant | WT | L258E | L258K | N262D | N262Y | K272E | S275H | S275R | N276G |
| --- | --- | --- | --- | --- | --- | --- | --- | --- | --- |
| IC<sub>50</sub><br>(ng/mL) | 290.4 | 1101 | 1890 | 1081 | 223.5 | >10000 | 3966 | 1633 | 756.4 |
| Fold<br>change | - | 3.8 | 6.5 | 3.7 | 0.8 | - | 13.7 | 5.6 | 2.6 |

**Figure 3.**
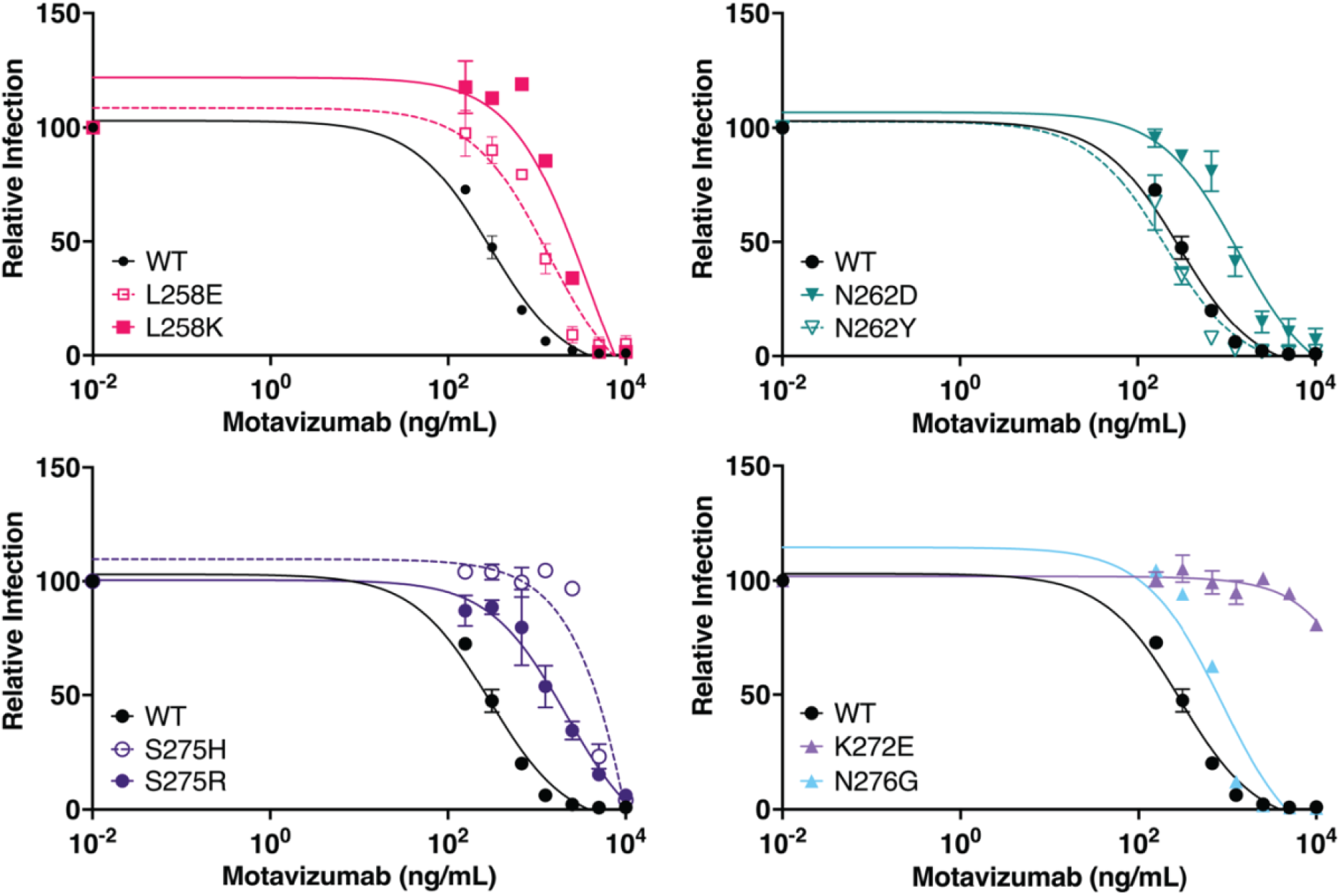
Microneutralization assay revealed reduced neutralization by motavizumab for six of the eight variants. Motavizumab was serially diluted two-fold (10000 ng/mL-156 ng/mL) and incubated with variants prior to infection of HEp-2 cells at an MOI of 1. mKate2-positive cells were counted by flow cytometry at 18 hours post-infection. Error bars indicate SEM of three independent experiments.

### MARMs had reduced motavizumab binding to F protein

Flow cytometry was used to assess motavizumab/F binding in a high-throughput format. HEK 293A cells were transiently transfected with plasmids that express variant F proteins. A subset of the variants that we had tested in the infectious clone for neutralization (L258E, L258K, K272E, S275H, S275R, and N276G) were tested as well as additional variants at the same residues of interest (L258V, K272G, K272R, and S275A). All of the additional variants tested, except K272G, had ΔΔ*G*_Bind_ values near 0 kcal/mol (Appendix) and thus would not be expected to have reduced binding by motavizumab. Expression of F protein was analyzed using mAb 101F that binds a distinct antigenic site from motavizumab (24). Motavizumab was conjugated with Alexa Fluor 488 (motavizumab-488) and 101F was conjugated with Alexa Fluor 594 (101F-594). Binding by 101F was used to gate on the F protein positive cell population and the median fluorescent intensity (MFI) of Alexa Fluor 488 was used as an indicator of motavizumab binding (Figure 4A). A reduction in binding was observed in the peak shifts and reduction of motavizumab positive cell percentages for L258E, L258K, K272E, K272G, S275H and S275R (Figure 4A). A significant reduction of MFI was observed in cells expressing L258E, L258K, K272E, S275H, and S275R when compared to WT (Figure 4B). L258V, N276G, and K272G had some reduction in MFI when compared to WT but were not statistically significant. K272R and S275A had MFI values similar to WT indicating that the binding of motavizumab is similar to WT, as expected because they had predicted ΔΔ*G*_Bind_ values of less than zero. All the MARMs that demonstrated reduced neutralization (Figure 3) also had reduced binding of motavizumab (Figure 4). There was a reasonable correlation (R^2^ = 0.52) when comparing binding of motavizumab to predicted ΔΔ*G*_Bind_ values, as ΔΔ*G*_Bind_ increases there is a decrease in the binding of motavizumab (Figure 4C), suggesting that an increase in ΔΔ*G*_Bind_ value may be more disruptive to antibody binding.

**Figure 4.**
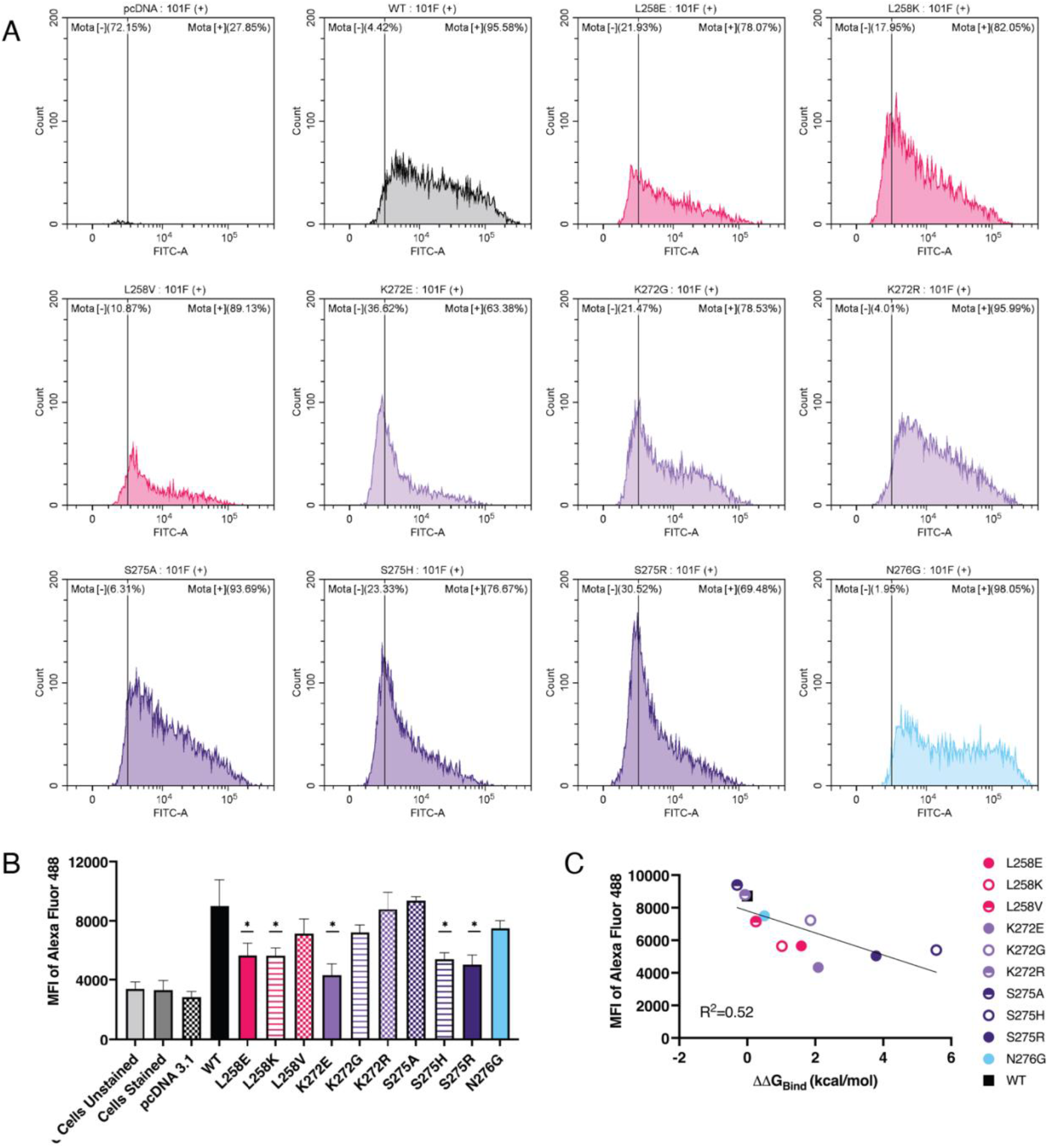
Flow cytometry revealed reduced binding of motavizumab to MARMs. Cells were transiently transfected with variant F protein-expressing plasmids or an empty vector control (pcDNA3.1) and dual stained with motavizumab-488 and 101F-594. Cells were gated on the 101F positive cell population, and the MFI of the motavizumab-Alexa Fluor 488 was quantified. (A) Histograms of motavizumab-488 positive cells gated on 101F-594 positive cell population are representative of three replicate experiments. (B) MFI of motavizumab-488 channel on 101F-594-gated cell population showing averages and SEM from three replicate experiments. \**p* <0.05. (C) Graph of ΔΔ*G*_Bind_ vs. average MFI of motavizumab-488.

### Escape from neutralization corresponds with altered on rate of motavizumab binding to F protein

To further evaluate motavizumab binding to MARMs we analyzed binding kinetics of the variants we identified to have the highest increase in IC_50_ compared to WT: K272E, S275H, and L258K. Surface plasmon resonance (SPR) was used to quantify binding kinetics of purified virions to motavizumab. A decrease in the binding on-rate (k_a_) was observed for all three variants, notably a >6-fold reduction of binding for S275H (Figure 5A, Table 3). There was no significant increase in the off-rate (k_d_) of antibody binding to the three variants (Figure 5B, Table 3). The overall equilibrium disassociation constant (KD) for the variants demonstrates a lower affinity between the MARMs and motavizumab when compared to WT (Figure 5C, Table 3).

**Table 3.** Binding kinetics of selected variants to motavizumab.

| Variant | $k_a$<br>(1/(M*s)) | Fold<br>Change | $k_d$ (1/s) | Fold<br>change | KD (M) | Fold<br>Change |
| --- | --- | --- | --- | --- | --- | --- |
| WT | 6.01E+05 |  | 4.64E-04 |  | 9.2E-10 |  |
| L258K | 2.26E+05 | -2.7 | 2.14E-04 | +0.46 | 9.56E-10 | -1.0 |
| K272E | 1.70E+05 | -3.5 | 2.32E-04 | +0.50 | 1.35E-09 | -1.5 |
| S275H | 9.81E+04 | -6.1 | 3.14E-04 | +0.68 | 3.01E-09 | -3.3 |

**Figure 5.**
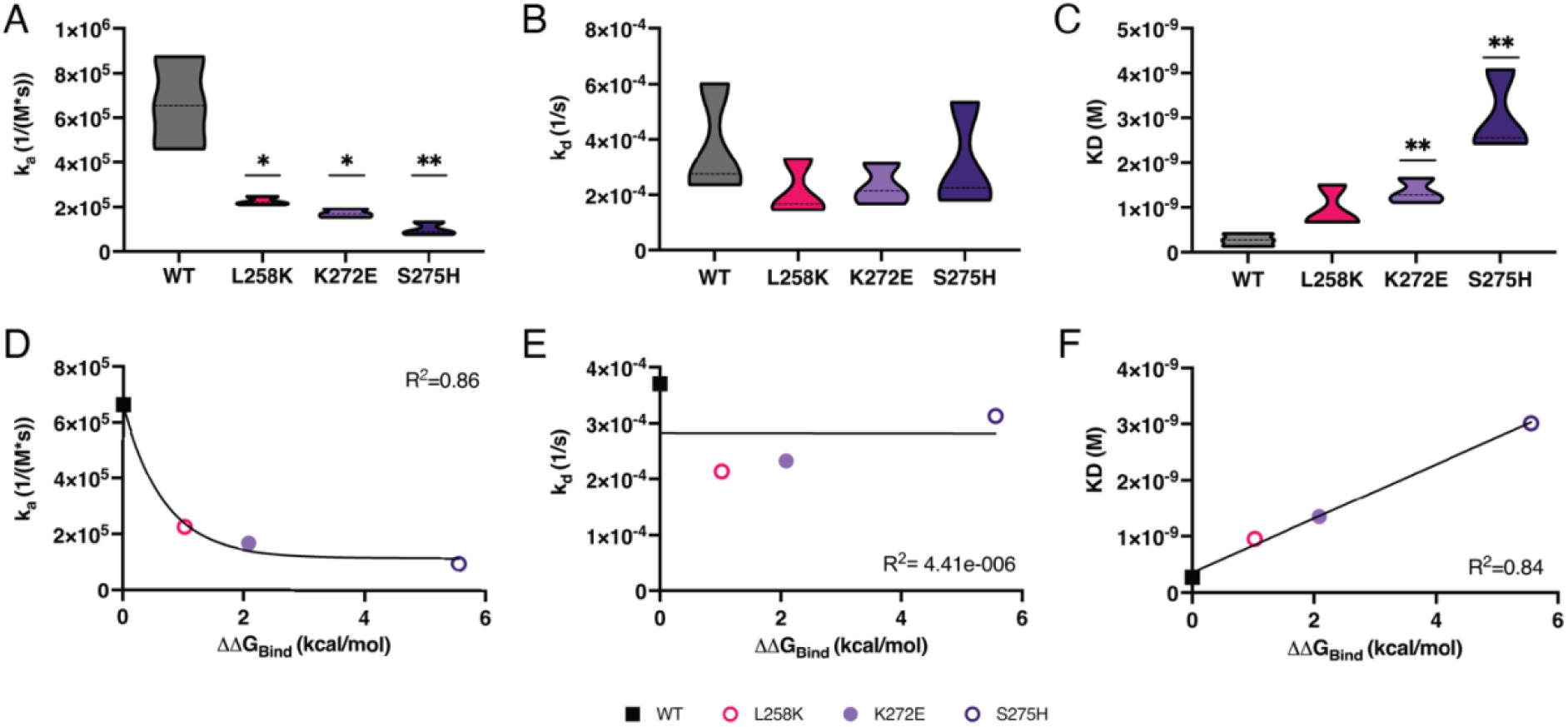
SPR revealed reduced on-rate as escape mechanism for variants. Purified virions were flowed over motavizumab at 40 nM, 20 nM, 10 nM, and 5 nM and (A) on-rate, (B) off-rate, and (C) affinity measurements were calculated. Three independent experiments were carried out for each variant. \**p* <0.05; \*\**p*<0.005; \*\*\**p*<0.001. Graphs of (D) on-rate, (E) off-rate, and (F) affinity and ΔΔ*G*_Bind_.

The low KD for WT is expected as once the antibody binds it is unlikely to be disrupted. There was a correlation between ΔΔ*G*_Bind_ and on-rate, as ΔΔ*G*_Bind_ increased the on-rate decreased (Figure 5D). No correlation was observed between ΔΔ*G*_Bind_ and off-rate, which is expected as there were no significant differences between the off-rate values (Figure 5E). A strong linear correlation was observed between ΔΔ*G*_Bind_ and overall affinity of motavizumab binding, as ΔΔ*G*_Bind_ increased, KD increased (Figure 5F). The escape mechanism for all three variants appears to be a decrease in association with motavizumab rather than an increase of disassociation from motavizumab.

## Discussion

The MD+FoldX approach provides rapid and testable predictions of MARMs. In this study, we applied this modeling approach to identify MARMs of RSV F protein and motavizumab antibody complex. This also enabled us to empirically validate the accuracy of our modeling approach. We selected eight variants that were predicted to allow the F protein to fold correctly but disrupt its ability to bind to motavizumab. All the selected variants were found to infect cells and propagate new virus. Six of the eight mutations demonstrated reduced neutralization by motavizumab when compared to WT and flow cytometry confirmed reduced binding of motavizumab to the mutants. SPR analysis of the variants that had the highest resistance to neutralization suggested that a reduced on-rate of antibody binding to virions was responsible for resistance and correlated with modeling estimations.

Previous studies have identified palivizumab and motavizumab MARMs from patient samples and passage experiments (19–27) and K272E was previously the only known MARM for motavizumab (21). This study examined mutations for motavizumab given the availability of the co-crystal structure and revealed novel escape variants. The binding kinetics of K272E have been previously studied using purified F protein and found that binding on rate rather than off rate appeared to be the mechanism of escape (27). In our study we were able to replicate those findings with purified virions as well as establish the same mechanism for escape for two additional MARMs that we identified. Previous studies have also found that antibody association rates correlate well with viral neutralization (37).

A limitation of the MD+FoldX approach and for molecular modeling in general is the reliance on having a co-crystal structure of the mAb and viral protein. Motavizumab is not used clinically but there are no co-crystal structures available for palivizumab and F protein.

However, most mAb in the pipeline for medical use have co-crystal structure with proteins of interest available (49, 50). Another drawback is that many of the viral proteins used in the crystal structures are lab strains of virus and may not reflect circulating viral strains. The RSV-A2 strain used in our study is based on a clinical isolate from 1961(51) and is the prototypic RSV strain for lab use. The F protein used for crystallography, DS-Cav1, is based off the A2 strain with modifications for expression and trimerization (52). However, the predominant circulating RSV A strain, ON1, was first identified in 2006 in Ontario and was classified as a novel genotype based off both F gene and G gene sequencing (53). The use of strains that reflect current circulating strains will help produce data that are more clinically relevant and can be done in the same reverse genetics system used for these experiments (54). Molecular modeling is also limited to predictions at individual amino acid residues. Studies of MARMs derived from a different F protein mAb, nirsevimab, have demonstrated that some MARMs require two separate residue changes for mAb escape (55). As more advances are made in fast protein-protein binding affinity estimation methods, we may be able to predict the effects of epistatic mutations.

In our previous study, comparing performances of different protein-protein binding affinity prediction methods, MD+FoldX approach showed the best performance for antigen-mAb complexes (56). It achieved an r value of 0.39 for 253 mutations from eight different antigen-mAb complexes with known experimental binding affinity data. We observed a slightly better performance of MD+FoldX approach for our smaller set of mutations tested in this study, where we obtained R^2^ of 0.52 between predicted ΔΔ*G*_Bind_ values and binding measured using flow cytometry (Figure 4C). There were strong correlations between the binding kinetics values and estimated ΔΔ*G*_Bind_, with R^2^ values for on-rate and affinity of 0.86 and 0.84, respectively.

However, this data set has only four data points and should be investigated further with more variants. Although this modeling approach remarkably identified six MARMs, there is still a need for better modeling strategies to improve correlation between experimental and predicted ΔΔ*G*_Bind_ values for antigen-mAb complexes. Use of MD simulation to provide conformation sampling is the key reason behind the performance of the MD+FoldX approach (56). However, empirical validation of the MD+FoldX approach carried out in this study further emphasizes the need for improved conformational strategies for large protein-protein complexes.

K272E was the only previously known MARM for motavizumab and no other study with passage experiments has been able to evolve other mutations. Future studies may investigate the evolutionary pathway that results in a poorly replicating MARM such as K272E becoming the dominant variant over more fit MARM. We can predict and generate MARMs, but further understanding of viral evolution is essential to refining our predictions and variant watch lists. Expansion of MD simulation and testing for other mAb such as nirsevimab and other viruses such SARS-CoV-2 and the respective mAbs would bolster our proof of concept. This would also be of interest and significance as MARMs have been identified for nirsevimab and both Delta and Omicron variants of SARS-CoV-2 have demonstrated resistance to therapeutic mAb treatments (55, 57, 58). In addition, mutations outside of the receptor binding domain of the SARS-CoV-2 spike protein have demonstrated increased instability in binding of mAb to MARM variants and would be of future interest for modeling studies (59).

In summary, biophysical protein modeling is a powerful tool for predicting MARMs. The ability to predict MARMs for current and future mAb treatments expedites variant discovery and has the potential to help design and optimize mAb. Given the rapid evolution of some viruses, such as SARS-CoV-2 and HIV-1, having a list of potential escape mutations is not only important for public health, but also for the scientific community to develop improved treatments for MARMs.

## Materials and Methods

### *In silico* predictions of MARMs using MD+FoldX approach

To predict RSV F protein escape mutations against motavizumab mAb, we applied our approach from previous studies that combines classical molecular dynamics (MD) and FoldX software (MD+FoldX) (15, 16). To designate a mutation as an escape mutation it requires: 1) disrupt binding to a mAb, and 2) leave the F protein monomer stable thus allowing it to fold and assemble. It is thus necessary to determine how amino acid mutations alter stabilities (ΔΔ*G* values) for F protein monomer folding (ΔΔ*G*_Fold_) and binding to Motavizumab (ΔΔ*G*_Bind_). Therefore, we used our MD+FoldX approach to estimate the folding stability of F protein monomer and F protein trimer/Motavizumab complex binding affinities due to all possible single mutations.

### Structure preparation

The X-ray crystal structure of RSV F glycoprotein bound to motavizumab was downloaded from Protein Data Bank. (PDB ID:4ZYP) 3D coordinates file was first modified to remove all but F protein trimer and three copies of heavy and light chains of motavizumab bound to each F protein monomer (44). The MODELLER software was then used to alter engineered residues and build the missing residues in all the chains (60). Missing amino acid residues 96 to 137 in F protein monomer represent liberated glycopeptide because of proteolysis by furin like proteases. These residues were ignored in our simulations.

### Molecular dynamics simulations

F protein monomer and F protein trimer/motavizumab complex structures were used as starting structures for MD simulations. Similar MD simulation the protocol was applied as reported in our previous studies for both the structures (15, 16). Briefly, the AMBER99SB*-ILDNP forcefield and the GROMACS 5.1.2 software package were used for generating topology files and performing simulation (61, 62). The final production simulation was run for 50 ns and snapshots were saved every 1 ns resulting in 50 snapshots for the F protein monomer and F protein trimer/motavizumab complex.

### FoldX

FoldX software was used to analyze MD snapshots of F protein monomer to estimate ΔΔ*G*_Fold_ and F protein trimer/Motavizumab snapshots were used to estimate ΔΔ*G*_Bind_ for all possible mutations at each site in F protein. Our FoldX analysis protocol involved processing each snapshot six times in succession using RepairPDB command to energy minimize the snapshot, BuildModel command to generate all possible 19 single mutations at each site in F protein, and then the folding (Δ*G*_Fold_) and binding affinity (Δ*G*_Bind_) were estimated using Stability and AnalyseComplex commands, respectively. Both folding and binding ΔΔ*G* values for each mutation was calculated by taking a difference between mutated and WT Δ*G*_Bind_ and Δ*G*_Fold_ values. For each mutation, we then averaged ΔΔ*G*_Bind_ and ΔΔ*G*_Fold_ values across all individual snapshot estimates. To estimate ΔΔ*G*_Bind_ values for all possible 19 mutations at each amino acid site of F protein, we performed 1,276,800 FoldX calculations (448 F protein residues × 19 possible mutations at each site × 50 MD snapshots × 3 copies of F protein/Motavizumab). Similarly, to estimate ΔΔ*G*_Fold_ values for all possible 19 mutations at each amino acid site of F protein monomer we performed 425,600 FoldX calculations (448 F protein residues × 19 possible mutations at each site × 50 MD snapshots). Averaging estimates across all individual snapshots ultimately resulted in 8512 ΔΔ*G*_Bind_ and ΔΔ*G*_Fold_ values for all possible mutations of F protein (see Appendix).

### Cell Lines

HEp-2 cells (ATCC CCL 23) were maintained in minimal essential media (MEM) with Earle’s salts, 10% fetal bovine serum (FBS), 1% MEM non-essential amino acids, 1% penicillin-streptomycin-amphotericin (PSF), and 5mM L-glutamine. BHK-21 BSR-T7/5 (63)were supplied by Dr. Ursula Buchhoz (NIH) and maintained in Glasgow’s MEM supplemented with 10% FBS, 2% MEM amino acids, 2mM L-glutamine, and 1% PSF. BSR-T7/5 were passaged every other passage with 1mg/mL geneticin to maintain the T7 polymerase-expressing plasmid. HEK 293 A cells were supplied by Dr. Lee Fortunato (University of Idaho) and were maintained in Dulbecco’s Modified Eagle’s media with 10% FBS, 1% penicillin and streptomycin, and 2 mM L-glutamine All cells were incubated at 37°C and 5% CO_2_. FreeStyle™ CHO-S™ Cells (Invitrogen) were maintained in FreeStyle™ CHO™ Expression Medium with 8 mM L-glutamine and incubated shaking at 37°C and 8% CO_2_

### Plasmid Preparation and Viral Propagation

Bacterial artificial chromosome (BAC) containing the antigenomic cDNA of RSV-A2 mKate2 and four RSV helper plasmids (human codon bias optimized N,P,L, M2-1) were provided by Dr. Martin Moore (Emory University). RSV BAC and helper plasmids (WT) were transfected into BSRT7 BHK-21 cells (48). Media and cells were harvested, sonicated in an ice water bath 3 times at 30%, and centrifuged at 2000 rpm for 5 minutes at 4°C. Supernatant media was harvested, flash frozen and stored at −80°C. A working stock of the WT virus was generated by infecting HEp-2 cells for 1-hour rocking at 37°C and 5% CO_2_., adding complete media, monitoring cells for fluorescence and cytopathic effect (CPE), and was harvested as described above when fluorescence was detected throughout the flask. The F gene was subcloned from RSV BAC into pBluescript SK +. Site directed mutagenesis to generate the variants of interest was performed by Bioinnovatise Inc. Variant F genes were cloned back into genomic RSV BAC and transfected into BSRT7 BHK-21 cells as described above. Each mutant was plaque purified from HEp-2 cells (64) and confirmed with sequencing. Working stocks of the mutant F gene viruses were propagated as described above. All viruses were titrated by 50% tissue culture infectious dose (TCID_50_) on HEp-2 cells.

### Antibodies

Motavizumab and 101F antibodies and plasmids to produce these antibodies were provided by Dr. Jason McLellan (University of Texas, Austin) (41, 52). Plasmids expressing the heavy and light chains were co-transfected into FreeStyle™ CHO-S™ cells (Invitrogen) in serum free FreeStyle™ CHO™ Expression medium. Cell medium was harvested 6-8 days post-transfection and concentrated using a Vivaflow 200. The concentrated medium was purified using a HiTrap protein A column (GE Healthcare) as previously described (41, 52) and dialyzed into PBS.

### Growth Curves

HEp-2 cells were infected at a multiplicity of infection (MOI) of 0.1 in MEM. Cells were rocked for 1 hour at 37°C and 5% CO_2_. Cells were washed twice with PBS and the second wash was saved as the zero-hour time point. Complete media was added after aspiration of the PBS. Cell media was sampled at 6, 9, 12, 15, 18, 24, 36, and 48 hours. Viral titers were performed by TCID_50_ assay on HEp-2 cells.

### Microneutralization Assay

Assay was performed as previously described (65). Briefly, motavizumab was serially diluted two-fold starting at 10000 ng/mL and ending at 156.5 ng /mL. Variants were incubated with antibody for 1 hour prior to infecting HEp-2 cells at an MOI of 1. Cells were harvested 18 hours post-infection and counted via flow cytometry. Inhibition curves and IC_50_ values were calculated using GraphPad Prism v.9.

### F gene and Sequencing

RNA was isolated from virus stocks using Quick-RNA Viral Kit (Zymo Research). Viral cDNA was generated using SuperScript™ IV VILO™ (Thermo Fisher Scientific). The F gene was PCR amplified using primers RSV F 5’ (5’GCAAGGATTCCTTCGTGAC3’) and RSV F 3’ (5’CACACCACGCCAGTAG3’) and Phusion High-Fidelity DNA Polymerase (Thermo Fisher Scientific). Sanger sequencing was performed by Elim Biosciences.

### F Gene Library and Expression Vector

The F gene variant library was generated by Twist Bioscience, based on the codon-optimized F expressing plasmid, pHRSVFoptA2, provided by Dr. Mark Peeples (The Ohio State University) (66, 67). Each amino acid change at 17 residues within 5 Angstroms of the motavizumab binding site, based on the co-crystal structure (44), were incorporated into the variant library. F gene variants from the library were cloned into mammalian expression vector pcDNA3.1. Bacterial stocks were plated on LB-ampicillin plates and colonies were picked and sequence confirmed to isolate each variant. Plasmids for transfection were purified using PureLink™ HiPure Plasmid Midiprep Kit (Invitrogen).

### Flow Cytometry

HEK 293A cells were transiently transfected with pcDNA3.1-F plasmid variants or pcDNA3.1 as an empty vector control using Lipofectamine 3000™ reagents (Invitrogen). After 6 hours cell media was added to variants L258E, L258K, L258V, K272E, K272G, and K272R to increase protein expression. After 24 hours cells were lifted using Accutase and washed in PBS. Motavizumab and 101F were conjugated using Alexa Fluor™488 and 594 Antibody Labeling Kits (Invitrogen). Cells were stained with 1 μg/mL of Alexa Fluor 488 conjugated motavizumab and 0.7 μg/mL of Alexa Fluor 594 conjugated 101F antibody. Flow cytometry was performed using a Beckman Coulter Cytoflex-S and data were analyzed using CytExpert software.

### Viral Purification

WT, L258K, K272E, and S275H clonal virus stocks were used to infect HEp-2 cells. Uninfected HEp-2 cells were included as a control. Cells and supernatant media were harvested when mKate2 fluorescence was observed throughout cell culture flasks.

Harvested cells and media were flash frozen in liquid nitrogen and rapidly thawed in 37°C water bath. The cells and media were then sonicated in ice water bath at 30% for 30 seconds three times. Cell debris was pelleted at 2000 rpm, supernatant medium was removed and centrifuged again. The supernatant medium was loaded into Ultra-Clear 1 × 3.5 in. ultracentrifuge tubes (Beckman Coulter) with a 15% sucrose cushion in 10 mM Tris–HCl, pH 7.5, 100 Mm MgSO4 and 0.25 M sucrose. Samples were spun at 20000g for 2 hours at 4°C and pellets were resuspended in PBS. F protein concentration was determined using RSV-F ELISA Kit (SinoBiological).

### Surface Plasmon Resonance

All SPR experiments were performed on a Nicoya OpenSPR™ Rev 3 one channel and Protein A Sensors Kit (Nicoya) was used to attach 30 μg/mL of motavizumab in PBS to the chip. Purified virions at concentrations 40 nM, 20 nM, 10 nM, and 5 nM were allowed to associate for 5 minutes at 20 μL/minute and disassociate for 5 minutes at 20 μL/minute. Chips were regenerated between runs with 10 mM glycine pH 3 and motavizumab was reattached. All kinetics analysis was performed in TraceDrawer 1.9.

## Acknowledgments

This research was supported by the Institute for Modeling Collaboration and Innovation sponsored by the NIGMS under award number P20 GM104420 and National Science Foundation EPSCoR Track-II under award number OIA1736253. This research made use of the computational resources provided by the high-performance computing center at Idaho National Laboratory, which is supported by the Office of Nuclear Energy of the U.S. DOE and the Nuclear Science User Facilities under Contract No. DE-AC07-05ID14517. The authors would like to thank Mr. Kevin Hutchison for excellent technical support and Dr. Paul Rowley for invaluable discussions.

